# Direct measurements of active forces and material properties unveil the active mechanics of early embryogenesis

**DOI:** 10.1101/2025.05.11.653307

**Authors:** Arthur Michaut, Alexander Chamolly, Olinda Alegria-Prévot, Carole Phan, Francis Corson, Jérôme Gros

## Abstract

Despite progress in probing tissue mechanics, direct long-term measurements in live embryonic epithelia are lacking. This limits our understanding of amniote embryonic morphogenesis, which takes place over hours or days. Here, we introduce a combined technological and analytical framework to directly measure active forces and material properties in developing quail embryos, in a minimally invasive way. We find that the embryonic epithelium behaves elastically on short timescales with a modulus ∼ 2 nN · μm^−1^ but flows over longer timescales with a viscosity ∼ 4 μN · s · μm^−1^, and that both properties are under active regulation. Notably, we demonstrate that cell division is an active epithelial fluidizer, with division rates correlating with tissue fluidity. This fluidization is crucial for the emergence of the primary embryonic axis, which we show is shaped by a force ∼ 2 μN. Altogether, our approach delivers a complete and quantitative view of early embryonic mechanics, and opens new avenues for biomechanical studies in live embryos and tissues.

## Introduction

Although inherently biophysical, morphogenesis has been largely explored through cellular and molecular approaches. These studies, while they provide valuable insights, can only give qualitative indications on the causal mechanisms driving tissue deformation. The tissue deformations that ultimately shape the embryo are governed by mechanical forces that actively generated by its cells, and by its material properties, which set the magnitude and the timescale of the response[1]. Whereas force patterns can be deduced from the spatial distribution and temporal evolution of actin filaments and myosin motors within cells or across tissues, in the absence of specific readout or markers, tissue material properties (such as viscosity and elasticity) cannot be readily identified and require advanced biophysical techniques. Material properties can also be inferred through data-driven theoretical models [2–7]. However, in the absence of direct measurements, the inferred tissue stresses and material properties remain free parameters, making it difficult to ascertain the biological relevance of these theoretical frameworks. Obtaining direct *in vivo* measurements of forces and material properties is thus essential to elucidate the causal mechanisms of morphogenesis and understand how cellular behaviors and molecular regulators ultimately drive tissue deformation. Indeed, recent studies measuring tissue rheology have provided novel insights into morphogenetic processes, such as the importance of transitions between solid-like and fluid-like states [8, 9] or tissue softening [10]. However, measuring mechanical properties remains a considerable challenge[11–13]. In general, embryonic tissues are not amenable to classical rheological measurements, owing to their inaccessibility, geometry or fragility. For these reasons, several strategies have been devised to measure mechanical properties *ex vivo*, where dissected tissues have been aspirated[9, 14, 15], stretched[16–20], or indented[21, 22]. To obtain direct *in vivo* measurements, magnetic beads or droplets have been injected to be actuated within the bulk of tissues[10, 23], surface tissues have been compressed by Atomic Force Microscopy (AFM)[24], and full embryonic bodies have been stretched[25, 26].

Early avian embryos, with their large size, external development, and accessibility, are exceptionally well-suited for dynamic and biophysical studies. At the egg-laying stage, the embryo consists of a large, flat epithelial disk (∼3 mm in diameter, ∼50,000 cells) attached to the vitelline membrane only at its edge, resembling a suspended monolayer of about 20-30 μm thickness. Upon incubation, large-scale tissue flows, driven by the graded contraction of a supracellular actomyosin ring at the interface between the embryonic central region and the outer ring-like extraembryonic territory, shape the entire embryonic region. This process results in the formation of the primitive streak, a transient midline structure prefiguring the anteroposterior axis, where cells internalize to form the embryonic germ layers, in a process named gastrulation. We previously inferred that the morphogenesis of the early embryo and its primitive streak is enabled by the fluid-like response of the epiblast and suggested that cell rearrangements induced by cell division might regulate tissue viscosity [27]. However, only *in vivo* measurements of the embryo’s mechanics can unequivocally validate these hypotheses and mechanical assumptions, more generally.

Here, we present a novel cantilever-based micromanipulator system designed to measure the rheology of the early avian embryo. This system allows us to apply controlled in-plane forces for extended periods of time to the monolayer while simultaneously tracking tissue deformation at a cellular resolution, and to extract absolute values of the short-term elasticity and long-term viscosity of the tissue. Modulating the rate of cell division through drugs or changes in incubation temperature, we show that both elasticity and viscosity are actively regulated, and uncover an approximately linear relationship between the rate of cell division and tissue fluidity. Inserting the measured value of tissue viscosity into an analytical model of gastrulation predicts the force driving this process, which we experimentally confirm by imposing a stalling force at the contractile margin of the embryo. Our study provides the first comprehensive quantitative mechanical characterization of early embryogenesis—measuring both active forces and material properties.

## Results

### A motorized micromanipulator to probe tissue mechanics

To probe the mechanical properties of developing embryonic tissues, we devised a motorized micromanipulator to apply forces using a flexible cantilever. To image embryos while manipulating them, the device was built to be fitted on the stage of an inverted confocal microscope, allowing both large-scale and cell-resolved live imaging of manipulated embryos. The device is constructed by connecting a vertical flexible fiber to a piezo-electric stage (Fig. 1a, Fig. S1). The fiber is terminated by a fluorescent bead that orthogonally contacts the embryo, which is cultured *ex ovo* on a semi-solid culture medium (Fig. 1b-c). By coating the bead with Celltak, a non-specific tissue adhesive, we can establish a strong contact, allowing us to apply forces within the plane of the epithelium. In response to a controlled displacement of the piezo-electric stage along the *x* axis, the cantilever bends, and its deflection (∆*x*) is quantified by measuring the difference between the displacement imposed by the piezo motor and the actual displacement of the fluorescent bead, which is detected by confocal imaging. In the limit of small deflections with respect to the cantilever length, we can directly compute the applied in-plane force *F* = *k*∆*x*, where *k* is the calibrated stiffness of the cantilever. This device allows us to apply forces in the 0.1 - 2 μN range. The deformation of the embryonic tissue around the moving bead (Fig. 1d) is simultaneously imaged in transgenic embryos expressing fluorescent reporters (membrane bound eGFP (memGFP) or tdtomato:Myosin (myo), to reveal cell boundaries, and quantified using Particle Imaging Velocimetry (PIV), as previously described [27].

**Fig. 1.**
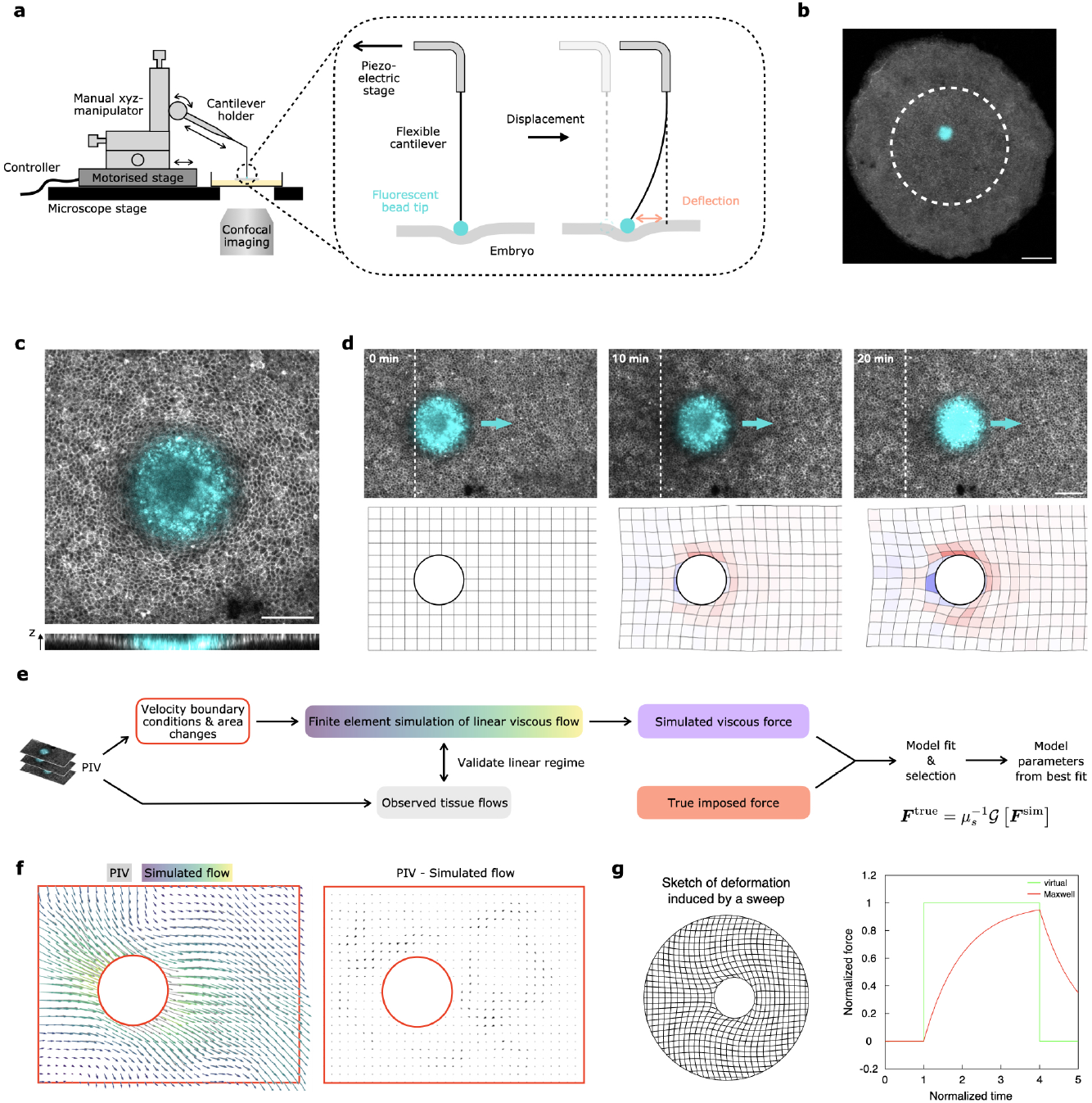
A cantilever-based micromanipulator to probe tissue mechanical properties. **a** Illustration of the micromanipulator system. **b** Low-resolution confocal image of an embryo indented with a bead (cyan) in its embryonic domain shown by a dotted line. **c** Cellular-resolution confocal image of the contact between the bead (cyan) and a memGFP embryo (gray), and *xz* orthoslice of the contact. **d** Top panels: time-lapse images showing tissue deformation driven by moving the bead towards the right (cyan arrow). Bottom panels: quantification of tissue deformation by PIV. **e** Flowchart of the model inference pipeline. Two-dimensional velocity fields are extracted from time-lapse images using PIV. Velocity values along an inner contour (surrounding the bead) and an outer contour (at the periphery of the imaging field) are used as boundary conditions for a finite element simulation of linear viscous flow in the enclosed domain. The output of this simulation is compared to the observed tissue flow to validate a description of the tissue as a uniform linear material. The time course of the simulated force exerted on the tissue from within the inner contour ***F*** ^sim^ is fit against the known true imposed force ***F*** ^true^ using a convolution with a parametrized viscoelastic model kernel *G* (see Methods). **f** Representative illustration of the finite element method. Left panel: boundaries of the mesh (red) superposed with PIV-derived flows (gray) and simulated flows (colored by magnitude). Right panel: boundaries of the mesh (red) superposed with the vector difference between the two flow fields (black). **g** Sketch of a deformation induced by a solid disk, and time courses of the virtual instantaneous force on the circle and the true force applied by a Maxwell fluid (time is normalized by the stress relaxation time, and forces by their steady-state value; with an imposed piece-wise constant velocity of the disk, the instantaneous force is also piece-wise constant). Scale bars: 500 μm (**b**), 100 μm (**c-d**)

### Computational inference of viscoelastic properties

To measure the mechanical properties of the embryonic tissue, the stresses imposed by the moving bead must be related to the tissue flows that these stresses generate. However, these flows are potentially affected by other factors such as the morphogenetic forces that shape the embryo, as well as the geometry of the contact with the bead and that of the embryonic disk. Although in this study we focus on the embryonic territory of the epiblast, which, as suggested by our previous work, is shaped through the passive transmission of forces generated at its margin, we note that even forces located far from the bead can generate long-range flows that affect the motion of the cantilever tip at a distance. As such, it is not possible to just infer a measurement from the recorded tip motion and cantilever deflection, but it is necessary to consider the two-dimensional flow pattern and stresses that surround the bead. While the tissue flows can be quantified accurately using PIV and the net imposed force is easily calculated from the cantilever deflection, the spatial distribution of stresses in the tissue is invisible and their computation already requires a hypothesis for a mechanical model. As such, iterating on the hypothesis to find a better match between model and experiment quickly becomes computationally expensive.

To circumvent this problem, we exploit the fact that the perturbations imposed by the cantilever are small and slow. As explained in Appendix B, this implies that the spatial pattern of motion in a domain surrounding the bead is fully determined by the instantaneous pattern of tissue expansion inside the domain and velocity at the boundary, independent of the shear response of the tissue. This holds even if the tissue has a history-dependent, viscoelastic response. It also implies that the instantaneous pattern of motion can be recovered using a purely viscous mechanical model that is simple to simulate, independently for each time step. From this, we can compute a virtual force on the bead that would be needed to drive the same instantaneous motion in a purely viscous fluid (see sketch in Fig. 1g). If we allow that the tissue actually has a viscoelastic response, then the time course of the force on the bead is fully determined by the time course of this virtual instantaneous force, independent of the spatial pattern of motion. This greatly simplifies the inference of tissue material properties, which is achieved by fitting the time course of the true force imposed by the cantilever ***F*** ^true^ against that of the virtual force ***F*** ^sim^ computed for a viscous fluid (Fig. 1e,f). This procedure assumes that no forces other than from the bead are generated within the simulation domain, which requires that there is no friction with a substrate, and that the measurement is carried out away from active actomyosin contractile sites. It does, however, account correctly for the cantilever deflection that is caused by the flows that such forces generate at a distance. The validity of this assumption can be checked visually and quantitatively by comparing the observed and simulated flow fields, and if necessary, the simulation domain can be adjusted to exclude regions that violate it.

The fit of ***F*** ^true^ against ***F*** ^sim^ is then carried out by convolving with a model-dependent kernel that is parameterized by mechanical properties such as viscosity and elastic compliance. Different hypotheses for viscoelastic models can be tested by comparing goodness-of-fit. Since ***F*** ^true^ and ***F*** ^sim^ only need to be computed once per dataset, fits can be rapidly fine-tuned and compared across a wide class of linear viscoelastic models.

### Sweeps on embryos under control conditions reveal Maxwell-like rheology

Using this framework, we set out to investigate the mechanical properties of the tissue at the long time scale (hours) over which gastrulation takes place. To this end, we performed linear sweeps across the epiblast with the stage moving at a constant 150 μm · h^−1^ for up to 2 hours, followed by a relaxation phase during which the stage remains stationary (Fig. 2a). The speed and duration of the cantilever’s displacement were chosen to drive tissue flows that are commensurate with the endogenous flows observed in unmanipulated embryos, and thus to apply a force that is physiologically relevant (Fig. 2b,c, Movie S1). During the sweep, the deflection built up steadily and partially relaxed afterward (Fig. 2d), as opposed to control experiments, where no contact was made with the embryo and the cantilever showed deflection (Fig. S2).

**Fig. 2.**
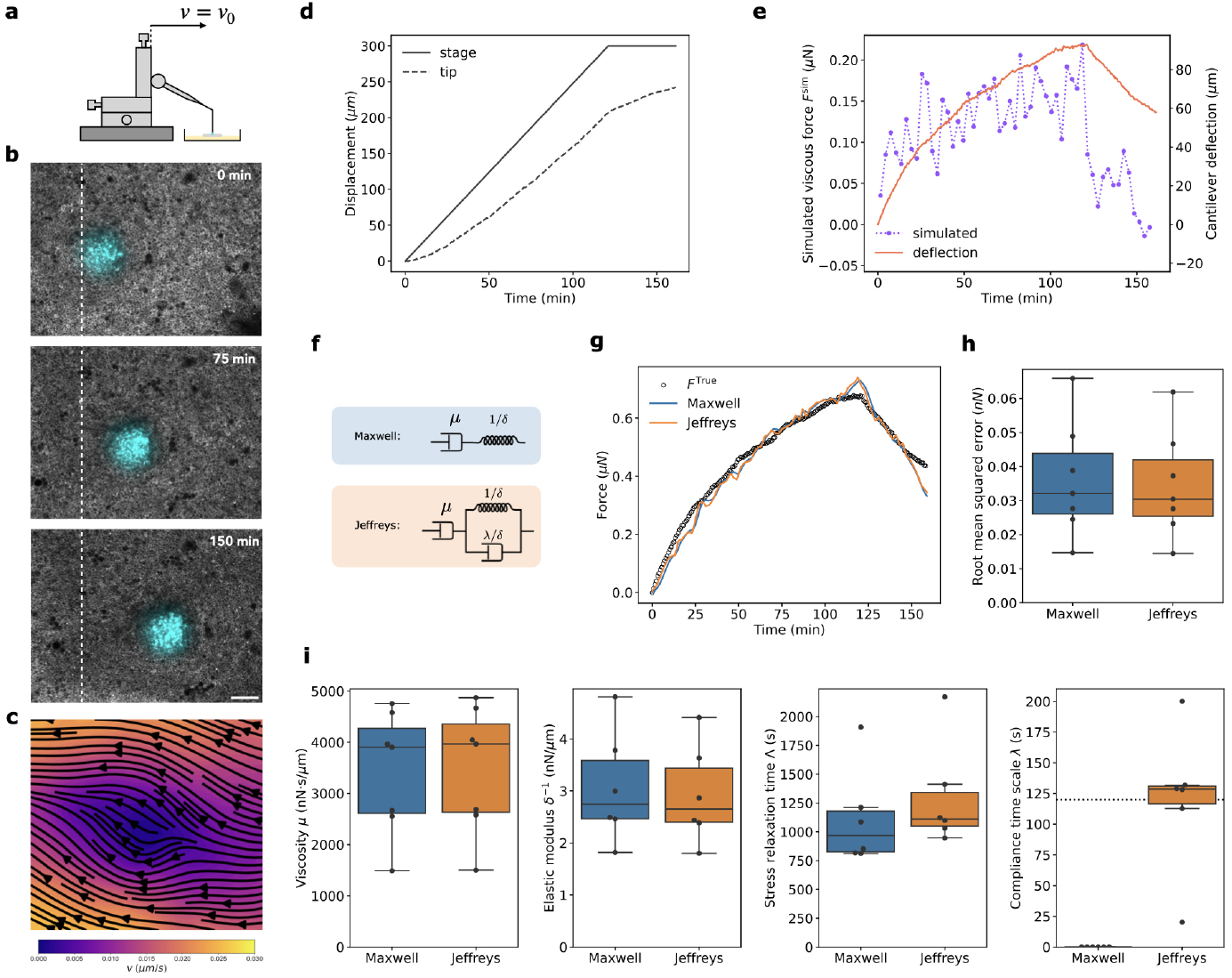
Long-time tissue viscosity measured by sweep assays. **a** Schematic of the imposed parameter: the stage velocity. **b** Time-laspe confocal images of a myo embryo (gray) with fluorescent bead (cyan). **c** Vvelocity field in a frame moving with the bead, computed by PIV at 75 min. **d** Recorded stage (solid line) and tip (dotted line) positions as a function of time. **e** Calculated cantilever deflection (red) and virtual instantaneous force (purple) against time. **f** Illustration of the models in terms of dashpot-spring networks. Maxwell’s model considers flow with viscosity *μ* with an additional elastic compliance of magnitude *δ*. Jeffrey’s model adds a non-zero time scale *λ* for the compliance. **g** True imposed force (black) and best fits of the different viscoelastic models from a convolution of the simulated force with the viscoelastic kernel. **h** Root mean squared error of the viscoelastic model fits (*N* = 7). **i** Fit parameter values for the viscosity (*N* = 7), elastic modulus and related time scales (*N* = 6) under control conditions. The dotted horizontal line indicates the time interval between frames used for PIV. Scale bar: 100 μm.

Using our finite element method, we can simulate the tissue response (Fig. 2e) to measure the mechanical properties using an appropriate mechanical model. We considered two potential models for the rheology: the Maxwell and Jeffreys models (Fig. 2f,g). The Maxwell model adds an instantaneous elastic compliance with modulus *δ* to a viscous flow with viscosity *μ*. This simple viscoelastic model notably features a non-zero time scale Λ = *δμ* on which stresses can be dissipated in a stationary configuration. In biological terms, this could be due to cell-cell rearrangements, which permanently change the configuration of the tissue. However, the Maxwell model predicts an instantaneous displacement jump in response to a step in applied force, which is a crude approximation. The Jeffreys model regularizes this feature by introducing an additional compliance time scale *λ* on which a steady flow state is reached. For this model, the stress relaxation time scale is modified to Λ = *δμ* + *λ*.

For *N* = 7 embryos under control conditions, we found the best fit to be provided by the Jeffreys model, closely followed by the Maxwell model(Fig. 2h). Considering the best fit for each dataset in more detail, we find very similar parameters for the viscosity *μ ∼*4000 nN · s · μm^−1^ and compliance modulus *δ∼* 0.35 μm · nN^−1^ for the Maxwell and Jeffreys models (Fig. 2i). The compliance time scale *λ* is null by definition for the Maxwell model, whereas it is fit to values around 125 s for the Jeffreys model. This value is comparable to the period at which ***F*** ^sim^ is sampled, suggesting that these sweep experiments could not capture precisely the short-term response of the tissue to loading.

### Step force assays at higher temporal resolution reveal constrained power-law response to stress loading

To explore further the viscoelastic behavior of the epiblast on short time scales, we next examined, with a higher time resolution (2 s per frame, vs 1 min per frame in the sweep assay) the response of the tissue to a step in applied force (in addition the higher time resolution, a step in force better probes the short-term response of the tissue that a sweep, which results in a progressive buildup of the force). To do so, we implemented a feedback loop to continuously adjust the stage position to match a prescribed force (Fig. 3a,b): after each stage displacement, the tip position, which is automatically detected in real-time, is used to compute the imposed force, and adjust the position of the stage at the next step. We used this method to impose a constant non-zero force, *F >* 0, for a duration of 1000 s followed by a relaxation step with a null force, *F* = 0. This assay allowed us to monitor the short-term response of the tissue (Fig. 3c,d, Movie S2), and extract the relevant mechanical properties using our finite element method (Fig. 3e,f). We considered again the Maxwell and Jeffreys models (Fig. 3g,h). At these time scales, the Jeffreys model considerably outperforms the Maxwell model (Fig. 3i), which, as noted above, predicts an instantaneous elastic displacement followed by steady viscous flow. Instead, the tissue exhibits a gradual transition between a fast initial response and a steady viscous flow, which is well captured by the Jeffreys model with a compliance time scale *λ*. However, we reasoned that this transient regime might exhibit a richer behavior than captured by a single compliance time scale. For this reason, we modified the Jeffreys model by replacing the short-time viscous element with a power-law element (Fig. 3f,g). This model yields the best fit, leading to a refined value of elasticity of 1.1 nN · μm^−1^, providing further evidence that embryonic tissues exhibit complex behavior at short time scales. Importantly, this step force assay and generalized Jeffreys model, while designed to better probe short-term properties, yield an average value of viscosity *μ∼* 5000 nN · s · μm^−1^ which is close to that obtained using the sweep assay and Jeffreys model. Altogether, these results demonstrate our ability to probe short-term elastic and long-term viscous properties of the developing embryo. Importantly, they show that over long time scales—most relevant to amniote embryo morphogenesis, which occurs over many hours (as for primitive streak formation)—the epiblast exhibits a viscous fluid response.

**Fig. 3.**
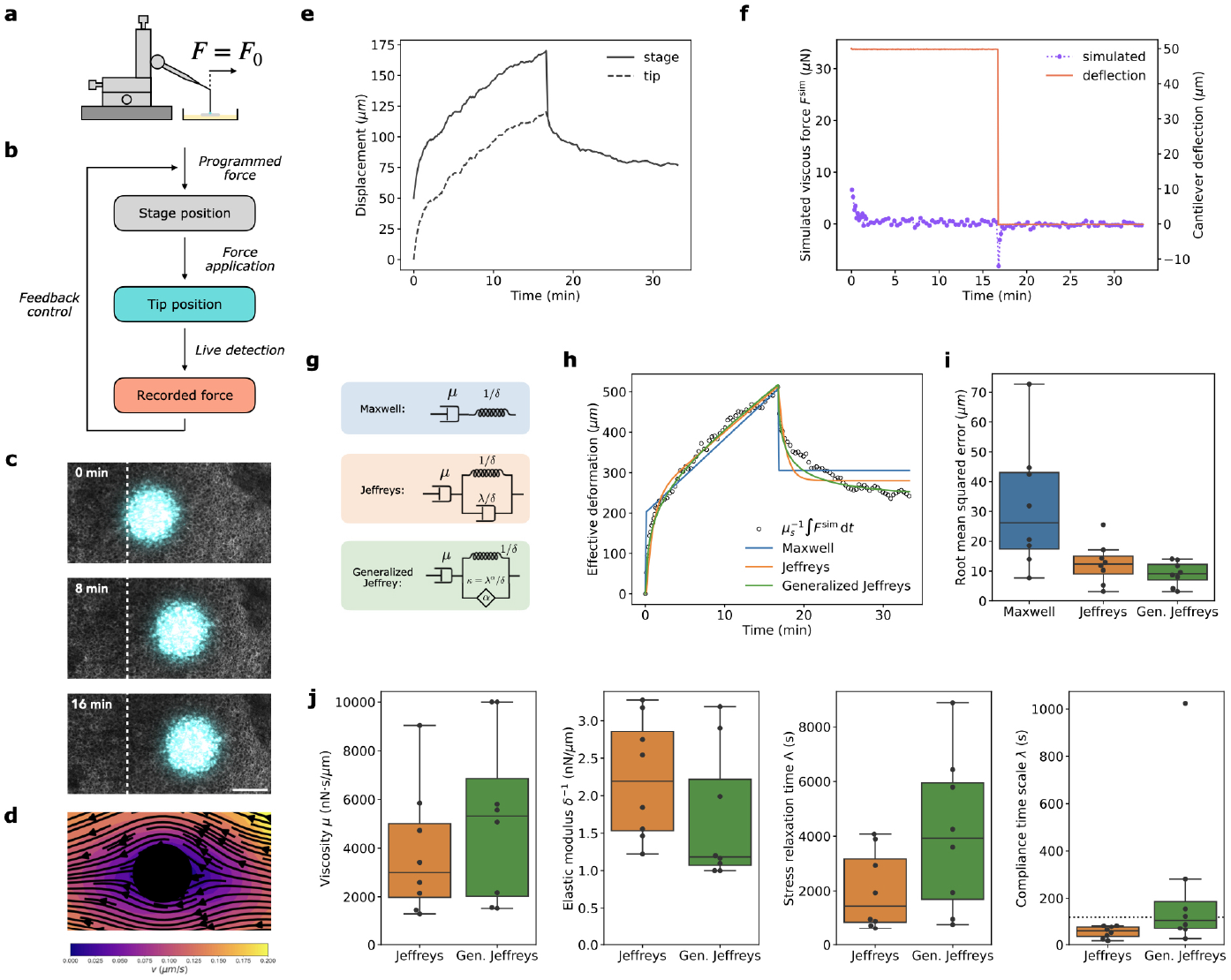
Short-time viscoelastic response measured by step force assay. **a** Schematic of the imposed parameter: the force. **b** Illustration of the feedback control loop to apply controlled forces. **c** Timelapse confocal images of a memGFP embryo (gray) with fluorescent bead (cyan). **d** Velocity field in a frame moving with the bead, computed by PIV at 1 min. **e** Recorded stage (solid line) and tip (dotted line) positions as a function of time. **f** Imposed cantilever deflection (red) and virtual instantaneous force (purple) against time. **g** Illustration of the models in terms of dashpot-spring networks. The Maxwell model considers flow with viscosity *μ* with an additional elastic compliance of magnitude *δ*. The Jeffreys model adds a non-zero time scale *λ* for the compliance. The generalized Jeffreys model with exponent *α* and modulus *κ* replaces the exponential compliance with a power law. **h** Time-integrated virtual instantaneous force (black) and best fits of three different viscoelastic models from a convolution of the true imposed force with a viscoelastic kernel. **i** Root mean squared error of the viscoelastic model fits (*N* = 7). **j** Fit parameter values under control conditions (*N* = 7). Where present, dashed horizontal line indicate bounds on the parameters. Scale bar: 100 μm.

### The epithelial embryo is actively fluidized by cell division

Our direct measurements of tissue mechanical properties support an effective description of embryo morphogenesis as the motion of a passive material driven by localized active forces. Nevertheless, we reasoned that embryonic tissues are active materials, and that their apparent material properties should be controlled by active cellular processes. In particular, lowering the incubation temperature, and thus reducing the rate of these active processes (cytoskeletal contractility and turn-over, cell division, etc.), should affect the material properties of the epiblast. We thus performed 2 h-sweep assays in embryos incubated at 21°C and 30°C, in addition to those performed at 38°C. As expected, morphogenetic movements were virtually arrested at 21°C, and were slowed down (compared to embryos incubated at 38°C) but were detectable at 30°C (Movie S3). Similarly, the tissue flows around the bead were progressively slowed down as temperature decreases, as revealed by examining tissue flows in a frame that moves with the bead (Fig. 4a). Using our finite element approach and the Jeffreys model to quantify the response, we found that, as temperature decreased, the elastic modulus of the tissue decreased, while viscosity increased, suggesting that the two properties (possibly through their dependence on cytoskeletal contractility and active cell rearrangements, respectively) are regulated by active biological processes (Fig. 4b). To gain insights into the cellular processes affected by temperature, we analyzed cell behaviors in the wake of the moving bead. Although acquired using a 10X objective, to capture a large tissue area around the bead necessary for the finite element-based analysis of the tissue response, groups of contiguous cells could be tracked in these movies (Movie S4). We found that at 21°C, the tissue is stretched by the moving bead, yet cells do not divide or rearrange, and for the most part, maintain their initial topology during the sweep assay (Fig. S3). This is in net contrast with control embryos incubated at 38°C, in which cells extensively divide and rearrange. We previously showed that cell division actively drives epithelial cell rearrangements of the early epiblast[27, 28], an effect that we proposed might regulate the tissue fluidity necessary to shape gastrulating embryos. To directly characterize the role of cell division in tissue fluidity, we first designed a segmentation-free automated pipeline for the detection of cell division and quantified their frequencies in the tissue surrounding the bead in embryos incubated at 21°C, 30°C and 38°C (Fig. 4c). Plotting these frequencies as a function of the measured fluidity (1*/μ*) from these same embryos revealed an approximately linear relationship between the frequency of division events and the measured fluidity (Fig. 4c). Second, to functionally test the role of cell division in controlling tissue fluidity, we inhibited cell division and measured the effect on mechanical properties. This was achieved using hydroxyurea (HU), in combination with Q-VD-OPh to prevent cell cycle arrest-induced apoptosis, as previously reported[27]. This treatment considerably slowed down both the endogenous morphogenetic tissue flows and the ones around the bead, resembling the ones quantified in embryos incubated at 21°C (Fig. 4a). We found that cell division inhibition led to a 4-fold increase in tissue long-time viscosity, similar to 21°C-incubated embryos (Fig. 4b). Consistent with a role for cell-division-induced rearrangements in the long-term behavior of the tissue, short-time elasticity was not affected by cell division inhibition. Altogether, these results demonstrate the role of cell division and the associated rearrangements in the control of tissue fluidity. In the embryo, tissue fluidity is thus an actively regulated property that is controlled by the rate of cell division.

**Fig. 4.**
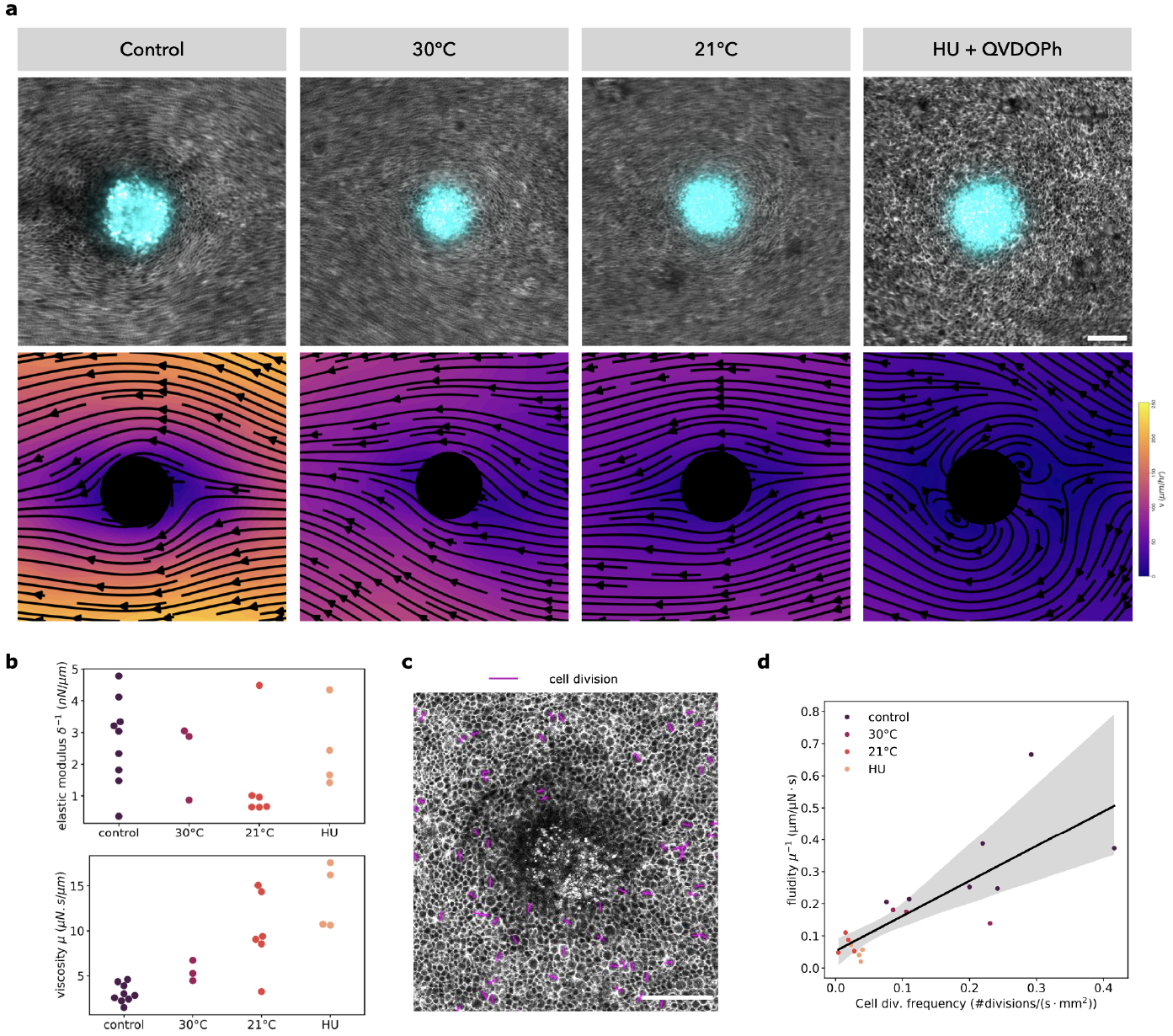
Impact of cell division on mechanical properties. **a** Impact of temperature and cell division inhibition by HU on the flow around the bead computed by registering the timelapse with respect to the moving bead. Top panels: time-averaged image over 10 min to show the structure of the flow. Bottom panels: streamlines computed by PIV. **b** Impact of temperature and cell division inhibition on the elastic modulus and viscosity. **c** Representative example of cell division automatically detected, the division main axis is indicated by a purple line. **d** Linear correlation between fluidity and cell division frequency. Scale bars: 100 μm

### A full quantitative mechanical description of early embryogenesis

Having determined an absolute average measure *μ* =4000 nN · s· μm^−1^ ≈1 μN · h· mm^−1^ for the epiblast viscosity, we were in a position to estimate the absolute force that shapes the embryo. To do so, we used a previously published model recapitulating gastrulation flows, which was based on the graded contraction of the margin and a viscous fluid response of the epiblast [27]. In the model, the pace of gastrulation is controlled by the difference ∆*T* in tension between the anterior and posterior margin, divided by the tissue viscosity *μ*. By fitting the model to experiments, we had estimated that this force/viscosity ratio is of the order of 2 mm · h^−1^ [27]. Our measurement of the tissue viscosity *μ* ≈1 μN · h· mm^−1^ now implies that the tension differential ∆*T* is of the order of 2 μN.

We further reasoned that this estimate could be verified by using the bead both as a probe and an obstacle to the flow along the lateral margin. The force exerted by the bead (through its link to the flexible cantilever) should increase gradually as it is displaced by the tissue, and eventually saturate when motion is stalled at the bead. To estimate the stalling force, we first considered a limit where the viscosity of the epiblast is small, such that forces generated at the margin propagate primarily along the margin (with a limited screening from the transmission of the force to the surrounding tissue [29]). In that limit, the force from the bead should leave relative motion along the margin unaffected, and induce a solid rotation of the embryo proper, overlaid on its endogenous motion (Fig. 5a,b). Under this simplifying assumption, the force needed to arrest motion at the bead is comparable in magnitude to the tension differential ∆*T* driving motion along the margin. Allowing for the fact that tension propagation along the margin has a finite range (commensurate with the distance between opposite sides [29]), the true stalling force should be a fraction of the driving force ∆*T*.

**Fig. 5.**
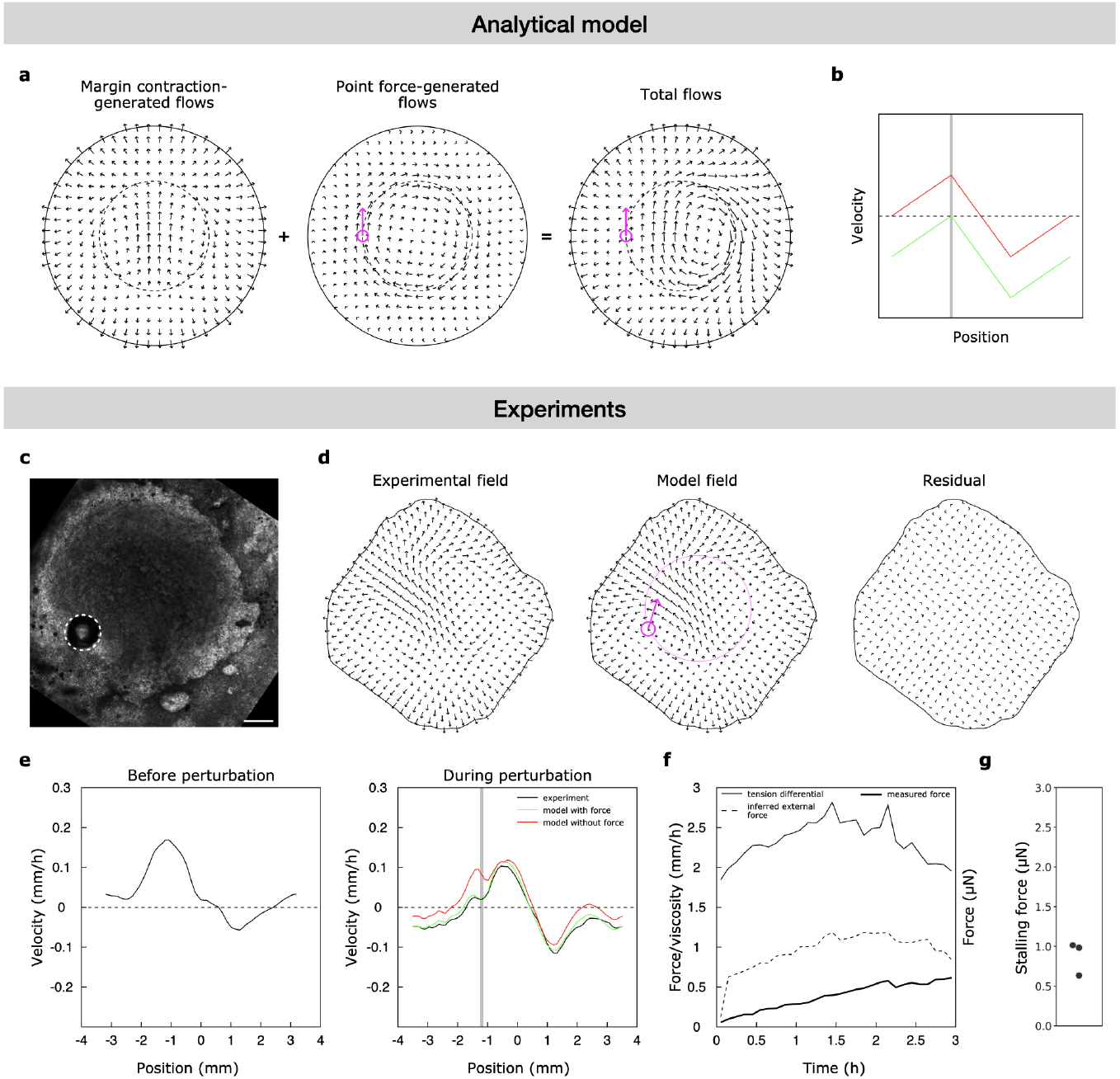
Measurement of the margin contraction force. **a-b** Analytical model of tissue movements with and without a point force applied to the margin (the force is assumed to propagate along the margin and drive a solid rotation). **a** Decomposition of the flow field of a gastrulating embryo with a point force applied to its margin. **b** Sketch of the tangential velocity along the margin without (red curve) and with (green curve) a point force. The gray line indicates the position of the bead. **c-g** Experimental measurements of the contraction force of the margin. **c** Image of a representative embryo with a bead indented in its margin. **d** Experimental flow field with the bead stalling the margin contraction, modeled flow field, and residual between the model and the experimental field. **e** Tangential velocity along the margin before and after the bead indentation (the latter compared with a model with or without a force on the bead). The gray line indicates the position of the bead. **f** Time courses of the tension differential and force on the bead inferred from the model, and of the measured force. **g** Measured stalling forces (maximum force reached when the bead stalls the contraction) for N=3 embryos. Scale bar: 500 μm

To test these predictions, we placed the cantilever-linked bead at the margin (Fig. 5c), at the estimated boundary of the posterior contracting domain of the margin (based on prior live imaging). Computing the tangential velocity along the margin, we found that, indeed, the bead skewed motion along the margin in the direction opposite to the initial velocity at the bead, but not as strongly as in the above idealization (compare Fig. 5b and 5e). Consistent with this, the absolute force on the bead plateaued under 1 μN (Fig. 5f,g), or about half of our estimate of the driving force ∆*T* ≈2 μN. To check that the ratio of about 1/2 between stalling and driving force is consistent with the actual pattern of motion in experiments, we modified our model of gastrulation flows [27] to incorporate the external force due to the bead. Based on fitting this extended model to experiments (Fig. 5d), we could infer force/viscosity ratios for the stalling force and the driving force in individual embryos (Fig. 5f). This recovered a ratio ∼1/2 between the two, which together with the direct physical measurement of the stalling force (Fig. 5f,g), corroborates our estimate of the driving force ∆*T* ≈ 2μN.

## Discussion

In this study, we present a bespoke yet readily replicable cantilever-based motorized micromanipulator mounted on an inverted microscope, which allows the minimally invasive application of controlled forces and deformations on live tissues while simultaneously monitoring their behavior. We furthermore establish a finite-element-based analysis framework that allows us to deduce absolute measures of short- and long-term mechanical properties in live embryonic epithelia from these mechanical assays. The system is versatile and can also be used to probe the active forces operating within developing tissues. Using this combined framework, we measured both the active force and viscoelastic properties underlying axis formation during early embryogenesis.

Similar approaches to probe tissue mechanics have been developed in early Drosophila embryos using magnetic beads or ferrofluid droplets [10, 30]. However, these studies probed only the short-term response of embryonic epithelia, isolated from morphogenetic events. In invertebrates such as Drosophila, morphogenesis proceeds rapidly, occurring over minutes to tens of minutes. By contrast, in amniote embryos, morphogenesis typically takes place over hours to tens of hours, and methods to probe tissues over longer time scales are essential to yield a physically relevant account of morphogenesis. In fact, we observed that the mechanical assays described here were gentle enough to allow embryos to continue to develop for hours and form an embryonic axis. Recently, a piezo-bender-actuated soft atomic force microscope (AFM) probe has been developed to probe the mechanics of early chick embryos, albeit at later stages and on short time scales [31]. However, in the absence of an accompanying framework to analyze tissue deformations, this could only provide relative indications on tissue material properties - and while useful for comparing different conditions or tissues, did not provide absolute measurements. Here, we provide a simple and computationally efficient approach to address this issue and obtain absolute values of viscoelastic properties from mechanical assays, even as the motion induced by the applied forces is overlaid on endogenous morphogenetic movements. This relies on two ‘tricks’. One, by simulating flows within suitably chosen regions of interest, the effect of ambient tissue motion is absorbed into the boundary conditions. Two, the spatial pattern of motion within the region of interest can be factored out, such that inferring tissue material properties is no more complicated than if a uniform deformation were applied to a tissue sample.

The measurements we obtain from our assays are two-dimensional, but can be extrapolated based on epithelial thickness (∼ 25 μm at this stage) to yield effective three-dimensional material properties, which are more intuitive to interpret. At 38°C, embryos exhibit a short-term elastic modulus of ∼100 Pa (comparable to soft hydrogels) and a long-term viscosity of ∼1.5 ×10^5^ Pa · s (similar to highly viscous fluids like putties or silicone greases). These values are consistent with those measured in other biological tissues (e.g. MDCK cellular aggregates [32] and chick presomitic mesoderm [9]). We show that both material properties are actively regulated, varying with temperature in a manner consistent with the modulation of active biological reactions, with elastic modulus and fluidity decreasing threefold at 21°C. That is, activity makes tissues stiffer against elastic deformations but more compliant to fluid-like flow. At 21°C, cells neither divide nor rearrange, in contrast with the numerous cell divisions and associated rearrangements observed at 38°C. Inhibition of cell division, which is the main driver of epithelial cell rearrangement at this stage [27, 28], led to a four-fold increase in epithelial viscosity. This result functionally validates the key role of cell division as an active epithelial fluidizer, which enables the irreversible tissue deformations underlying the emergence of the embryonic axis. The role of cell division as a fluidizer has been proposed both theoretically [2] and experimentally in the context of zebrafish gastrulation [15, 33]. However, these studies focused on mesenchymal tissues, and in the case of zebrafish gastrulation, division controls tissue viscosity by causing a transient loss of cell cohesion due to mitotic rounding. In contrast, epithelia remain cohesive even during cell division. Our work demonstrates that division-driven rearrangements are sufficient to fluidize the epithelial tissue without loss of cell cohesion. Importantly, inhibition of cell division did not significantly affect the short-term elasticity of the embryo. Although we do not directly address the biological basis of elasticity, it is reasonable to speculate that it is governed by components of the cell cytoskeleton, as shown in other systems [10, 30]. Recently, we showed that the early embryo exhibits an unusual behavior, flowing viscously under shear stresses but responding elastically to isotropic stresses on similar timescales [34]. Here, we designed sweep assays that probe the viscous response of the tissue to shearing. In the future, it will be of interest to design new mechanical assays that apply isotropic stresses to probe the long-term elasticity of the embryo, which is critical for shaping the early extraembryonic tissue.

## Acknowledgements

Work in J.G lab is funded by the Agence Nationale de la Recherche (EMBRYONICS ANR-19-CE13-0024 to J.G and F.C and LabEx Revive 10-LABX-0073), the European Research Council (ERC) under the European Union’s Horizon 2020 research and innovation programme (grant agreement n° 866186 to J.G), the Centre National de la Recherche Scientifique (CNRS) and Institut Pasteur. A.M is a recipient of a Pasteur-Roux-Cantarini postdoctoral fellowship. We would like to thank Albane Imbert, Eglantine Vignal and Eric Nicolau from the Institut Pasteur’s Fablab for their technical support. We also thank Thibaut Divoux for fruitful discussions. For the purpose of open access, the authors have applied a CC-BY public copyright license to any Author Manuscript version arising from this submission.

## Materials and Methods

### Animals

Animal husbandry for transgenic quails was carried out in accordance with the guidelines of the European Union 2010/63/UE, approved by the Institut Pasteur ethics committee authorization #dha210003, and under the GMO agreement #2432.

### Embryo imaging

Transgenic memGFP quail embryos were collected at stage XI using a paper filter ring and cultured on a semi-solid nutritive medium of thin chicken albumen, agarose (0.2%), glucose, and NaCl, as described in [35]. The embryos were then transferred to a bottom glass six-well plate (Mattek Inc.) with 2 mL (or 0.6 mL for high-resolution imaging) of the nutritive medium and imaged at 38°C (except for temperature experiments) with an inverted microscope Zeiss LSM 980 using a 2.5X/NA 0.12, or a 10X/NA 0.45 objective. The time interval between two consecutive frames was 1 min long-time sweep assays and 2 s for short-time long. Long-time sweep assays were monitored simultaneously with the 2.5X and 10X objectives by alternating the objectives between each frame.

### Cantilever fabrication and calibration

Taking inspiration from a suspended monolayer stretching device [36], the cantilever is made of a flexible fiber of 80 μm in diameter (Fort Wayne Metals, NitiWire) with a fluorescent bead of 250 μm glued at its extremity (Cospheric, UVPMS-BB-1.13 250-300um). The fiber was calibrated by measuring its deflection imaged with a stereoscope placed sideways. The deflection was generated by placing weights at the extremity of the fiber positioned horizontally. The force / deflection relation was linear and led to a fiber stiffness of 0.2378 N · m^−1^ measured by linear regression for a fiber of 14.5 mm in length. Using beam theory, the fiber stiffness can be computed for any given length (see Appendix A).

### Manipulator fabrication

The cantilever was manually positioned on the embryo using a manual XYZ manipulator (Thorlabs DT12XYZ/M). During experiments, the cantilever basis was actuated by a piezo-electric linear stage with a 500 μm traveling range (PhysksInstruments P-625.1CD). The cantilever was connected to the manipulator using a custom glass holder. A 15 cm long borosilicate capillary (outer diameter: 1 mm, inner diameter: 0.58 mm, Sutter Instruments) was pulled using a filament-based pipette puller (P-97 Sutter Instruments), the pulled capillary was severed to match the cantilever diameter. The cantilever end was inserted in the capillary and glued. The glass holder was then bent to the desired angle with a flame so the cantilever could be positioned vertically above the embryo. The glass holder was connected to the manipulator using a pipette holder (IM-H1, Narishige).

### Cantilever contact with the embryo

The cantilever bead is coated with Cell-Tak (Corning, 354240) for 1 hour prior to the experiment. It is then rinsed with PBS. Before putting the cantilever in contact with the embryo, the embryo and the imaging dish are covered with oil to prevent dehydration. Adding the oil layer is a delicate step that can detach the embryo from the vitelline membrane. For this reason, we optimize the oil viscosity by mixing a 3:1 ratio of mineral oil (Sigma) and silicon oil (Sigma). The cantilever is then positioned above the region to measure and manually lowered until contact is made. The pressure of the contact is gradually increased by inspecting in real-time the indentation of the monolayer by confocal imaging. The strength of the contact is then optimized by testing the absence of slippage during a short oscillatory solicitation.

### Cantilever tip detection

The cantilever tip is detected using a pattern recognition method. A reference image of the bead in contact with the embryo is defined before the experiment by manually cropping a region of interest around the bead. This image is then compared during the experiment (for real-time detection) or *a posteriori* (if real-time detection is not required) using a cross-correlation detection provided by the function skimage.feature.match_templatefrom the Python scikit-imagepackage [37].

### Detection calibration

To evaluate the precision of our setup, we calibrated our system with two kinds of assays. First, we run a linear sweep in the air to verify that the tip and the stage moved synchronously (Fig. S2a-b). Second, we run a similar sweep with the bead indented at the surface of a 452 Pa· s viscosity standard oil (Cannon). We verified that the displacement was linear and the measured force constant as expected for a pure viscous fluid (Fig. S2c-e).

### Force measurement

To measure the cantilever deflection, we measure the differential displacement between the cantilever tip and the cantilever basis which is connected to the stage. The stage position *x*_stage_ and the tip position *x*_tip_ are measured within two different frames of reference. *x*_stage_ stems from the stage’s physical position, while *x*_tip_ is measured by image processing. If the cantilever is not bent, the two positions vary concurrently when the stage is moved, which implies that the difference between the two positions *δ* = *x*_stage_ − *x*_tip_ stays constant. If the cantilever is bent, then the two positions don’t vary concurrently, which implies that *δ* varies. We can then compute the deflection ∆*x* by comparing *δ* to its value at rest *δ*_0_: ∆*x* = *δ* − *δ*_0_.

### Force controller

We developed a Python-based graphical interface to operate the stage and program mechanical assays. This interface, which is designed to control one (or several) stage(s), is distributed as a Python package named manipylator: https://pypi.org/project/manipylator/. manipylatorcan be used to maintain the applied force to a prescribed value in real-time. To do so, we employ a Proportional-Integral-Derivative (PID) algorithm provided by the Python package simple-pid(https://pypi.org/project/simple-pid/). This algorithm is controlled by 4 parameters: the 3 relative weights of the 3 PID terms and the frequency at which the correction is run. The correction frequency is limited by the image acquisition rate, as calculating corrections at a higher frequency than the image acquisition rate will lead to erroneous measurements of the deflection. We therefore typically set this frequency to the same order of magnitude than the acquisition rate. Regarding the PID weights, since the corrections are typically extremely small as the viscosity of the system makes it slow to react, a simple purely proportional feedback loop led to the most stable behavior.

### PIV and finite element modelling

Velocity fields were calculated from experimental images using a custom PIV software [27]. Velocity fields were computed on a square grid with 64 pixel resolution on images taken at time intervals of 120 seconds (sweeps) and variable intervals between 4 and 20 seconds (steps). Images were pre-processed with a convolutional length scale filter to improve detection of cellular motion. Tissue flows were simulated using the finite element python package FEniCSx v0.5.0 [38–41] using a velocity boundary condition derived from interpolating the PIV velocity fields linearly on boundary contours defined as the perimeter of the field of view (outer) and a 120 μm radius around the detected instantaneous centre of the bead (inner), adjusted in certain cases to account for irregularities in the image. Meshes were generated independently for each time step of the simulation, with a triangle size between 1*/*4 and 1*/*8 of the PIV and boundary data resolution. The quasi-steady nature of viscous flow allowed for the parallel and independent computation of the instantaneous force at each time step using P2/P1 Taylor-Hood elements for velocity and pressure. The flow was modelled as incompressible, with a prescribed divergence calculated from the PIV velocity fields, and the simulated force ***F*** ^sim^ was computed as the instantaneous stress exerted on the tissue by the inner contour. Net fluxes through and stresses on both the inner and outer contour were computed as error control metrics.

### Cell division inhibition

Cell division was inhibited by diluting 40 mM hydroxyurea (HU, Sigma) directly into the culture medium. To suppress the apoptotic effect of HU, we combined it with 1 mM Q-VD-OPh (Sigma, re-suspended in DMSO) as previously shown in [27]. HU is an inhibitor of the S phase; therefore, embryos were treated for 3 hours prior to imaging in order to let cells that had already passed the S phase complete their cell cycle.

## Supplementary figures

**Fig. S1.**
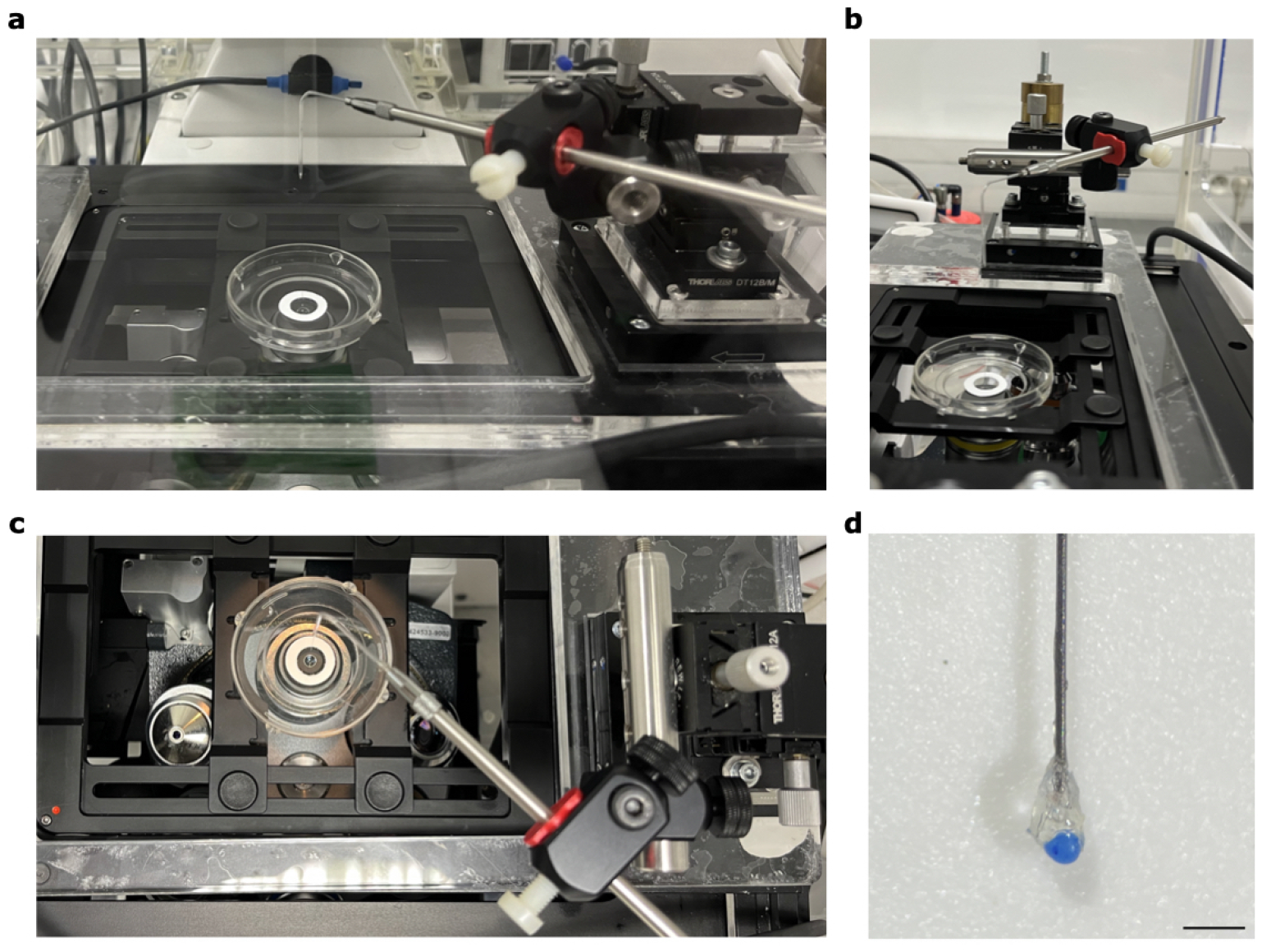
Experimental setup for manipulating avian epiblast. Front (**a**) side **(b**) and top (**c**) view of the setup. **d** Close up of the cantilever tip. Scale bar: 500 μm

**Fig. S2.**
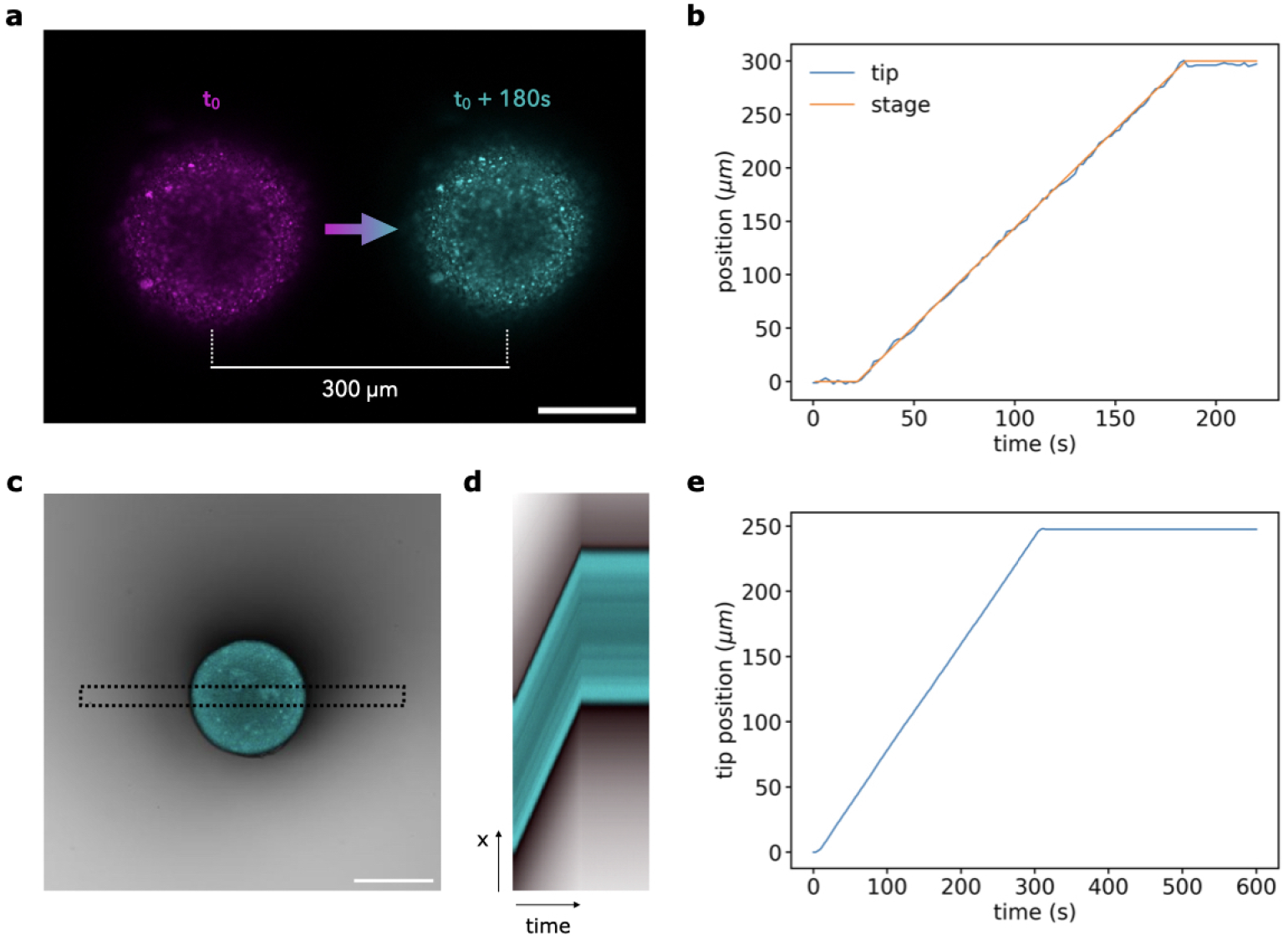
Evaluation of tip detection precision upon piezo actuations. **a-b** Control linear 300-*μ*m linear sweep without any contact with a substrate. **a** Color-coded starting (magenta) and final (cyan) positions of the tip during the sweep. **b** Synchronous displacement of both the tip and the motorized stage. **c-e** Linear sweep at the surface of a viscous oil. **c** Snapshot of the bead at the surface of the oil. The dotted rectangle indicates the region used to compute the kymograph in **d. d** Kymograph of the sweep. **e** Displacement of the tip. Scale bars: 100 μm

**Fig. S3.**
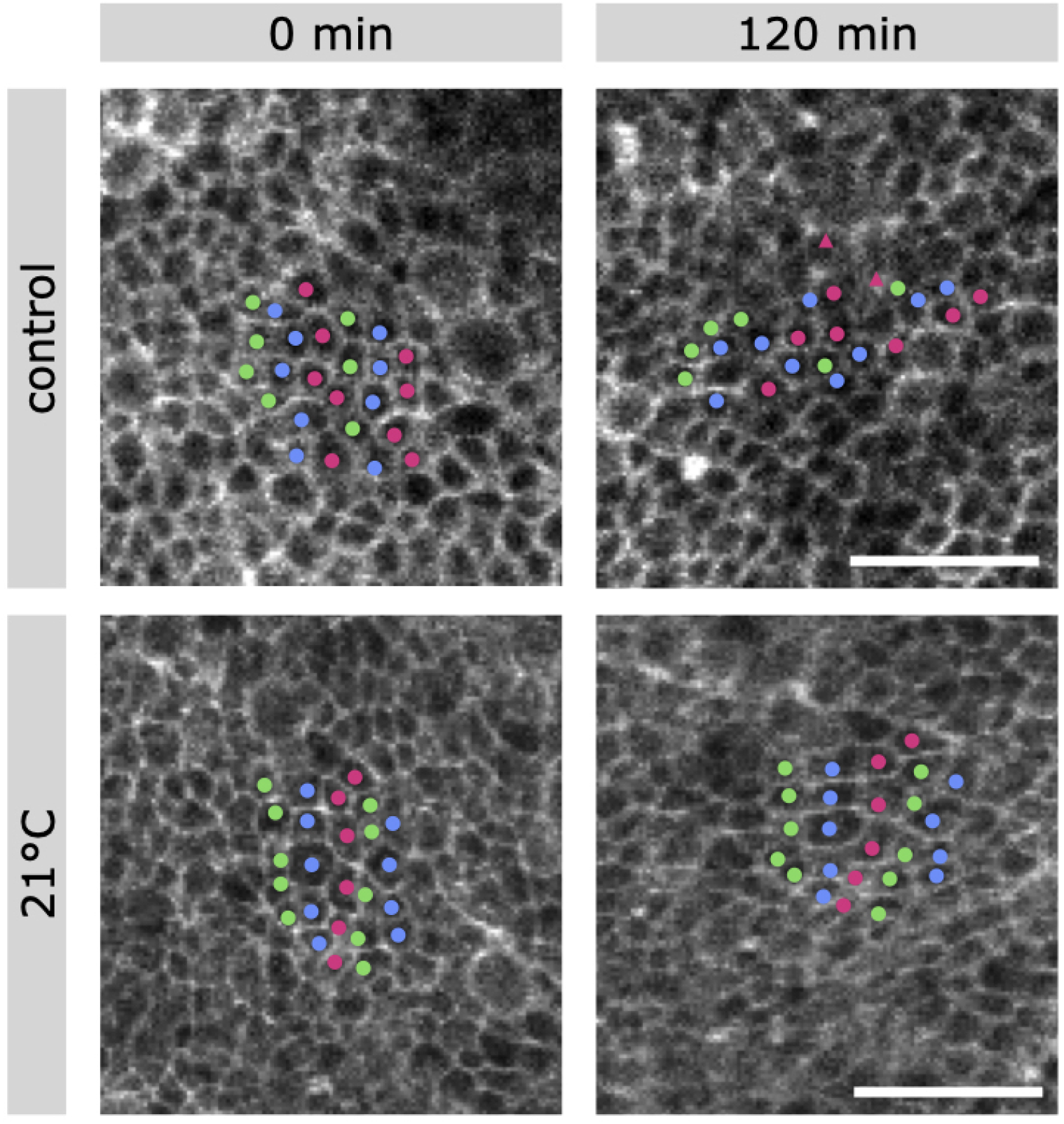
Cell rearrangements during a linear sweep in control condition and at 21°C. Cells were manually tracked and daughter cells are indicated by a triangle. Scale bars: 50 μm

## Supplementary movies captions

- **Movie S1**. Timelapse of a long-time linear sweep assay.
- **Movie S2**. Timelapse of a short-time force control sweep assay at a constant force of 365 nN followed by a relaxation phase at 0 nN.
- **Movie S3**. Effect of temperature and cell division inhibition during linear sweeps.
- **Movie S4**. Manual tracking of cell rearrangements in control condition and at 21°C.

## Appendix A Cantilever stiffness

According to Euler-Bernoulli beam theory, a light rod of length *L* that is clamped at one end and subject to a shear force *F* at the other has the profile [42]

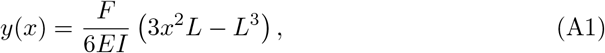

where *E* and *I* are the elastic modulus and moment of inertia respectively. Thus, the deflection at the free end is *y*(*L*) = *FL*^3^*/*3*EI*. It follows that the cantilever acts as Hookean spring with stiffness

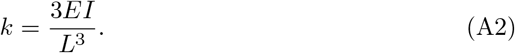

We measured the stiffness *k*_0_ of a reference cantilever with length *L*_0_. For different cantilevers with different length *L*, the stiffness *k* was calculated as 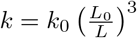.

## Appendix B Fitting procedure

Following [27], we focus on shear deformations, which shape the embryo proper, and do not seek to model the bulk response of the tissue. In modeling tissue flows within the embryonic disk, tissue area changes, including contributions from changes in apical cell areas (which contribute to a reversible stretching of extraembryonic tissue [34]), as well as cell proliferation or ingression, are incoroporated as a source term in the equations, by prescribing the divergence of the flow. Shear deformations, on the other hand, are described by a linear viscoelastic model, which relates the deviatoric stress tensor *τ*_*ij*_ and the shear rate tensor 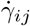 as

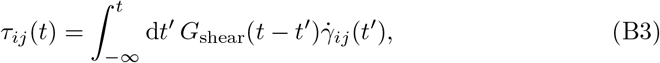

or using the operator notation for convolution with the viscoelastic kernel as described in section D.3

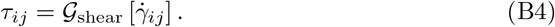

Here the shear rate tensor is defined in terms of the tissue flow vector ***u***(***x***, *t*) and the instantaneous area changes Γ(***x***, *t*) as 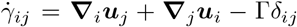, and we note that these equations are only valid as written in the small amplitude, due to the absence of non-linear convective derivatives [43]. As in the case of an incompressible fluid, the constraint of a prescribed divergence is enabled by a pressure *p* that contributes to the total stress tensor *σ*_*ij*_ together with the deviatoric stress as *σ*_*ij*_ = − *pδ*_*ij*_ + *τ*_*ij*_. Substituting into the Cauchy equation for the dynamics of an inertia-free material subject to a body force ***f***, **∇** · ***σ*** = −***f***, and dropping the subscript ‘shear’, we arrive at the constitutive equations

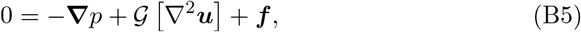

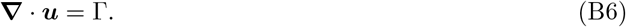

By applying the inverse operator 𝒢 ^−1^ to the top equation we see that there is a mapping between a Stokes flow with viscosity *μ*_*s*_ and a general linear viscoelastic flow:

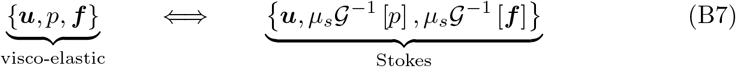

From this it follows that it is not possible to determine the rheology just from observing the kinematics – a different distribution of forces and pressure can give to the same observed flow for any linear and incompressible viscoelastic material. The mapping also carries over to the stress tensor:

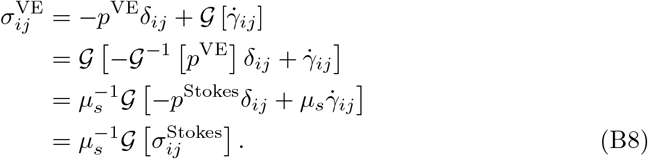

By integrating ***σ***^VE^ · ***n*** around a contour enclosing the point of contact between the cantilever and the tissue, we conclude the following relationship between the true force that is imposed on the tissue, ***F*** ^VE^, and the force ***F*** ^Stokes^ that is calculated from a finite element simulation of the observed flows assuming a linear viscous rheology with prescribed area changes and ***f*** = **0** in the simulated region:

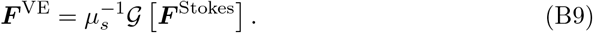

With this relation it is possible to fit a model to an experimental dataset. This fit can be carried out in two different directions, depending on whether ***F*** ^VE^ or ***F*** ^Stokes^ is considered the independent variable. For the other direction we write therefore:

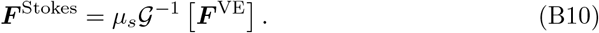

In practice, ***F*** ^VE^ is available at finely grained time points and is observed with little noise, while ***F*** ^Stokes^ is limited in time resolution by the PIV time step (usually 120 seconds), and noise both from the PIV and the accuracy of the finite elements. This noise can be partially mitigated by integrating ***F*** ^Stokes^ in time. Since the operator entails an integration, this is automatically provided by the fitting of ***F*** ^Stokes^ to ***F*** ^VE^ according to Eq. (B9). For the reverse case of fitting the imposed force to the simulated force, it is possible to integrate Eq. (B10) as

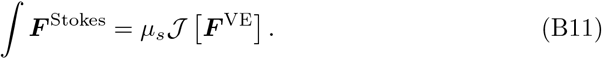

All we need then to fit a model to the data are the operators 𝒢, 𝒢^−1^ and 𝒥.

In practice, the 𝒢-fit described by Eq. (B9) yields the best results for linear sweeps, while the 𝒥-fit described by Eq. (B11) works best for step forces. Both fits are smooth, and for linear sweeps the 𝒢-fit benefits from a larger number of data points to fit against since ***F*** ^VE^ is resolved better than ***F*** ^Stokes^. For step forces, the advantage of the 𝒥-fit is mainly aesthetic, except that for the GJ model it is also easier to calculate numerically than the 𝒢-fit (see section C).

Since the 𝒢^−1^ operator is less smooth than either of the other two, Eq. (B10) is not the preferred choice to fit either type of data.

### Prestress

Occasionally, we would like to improve a 𝒢-fit to linear sweep data sets by allowing for a prestress in experiments that were not performed from an initially undisturbed configuration. We model this prestress phenomenologically as due to an arbitrary step strain just before the start of the experiment: 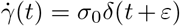 with *σ*_0_ the amplitude and *ε >* 0 the time interval between the perturbation and the start of the experiment. In this case, Eq. (B9) is modified to

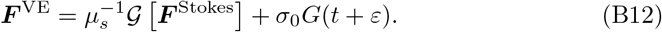

This introduces two additional fitting parameters, *σ*_0_ and *ε*. For the Maxwell and Jeffrey models, *G* is an exponentially decaying function and therefore the shift *ε* can be absorbed into the amplitude *σ*_0_. For the power-law model, this is not possible since the stress relaxation depends on the history of deformation: mathematically, *G*(*ε*) diverges as *ε →* 0 and in principle it is necessary to consider the delay *ε* as an independent parameter. In practice, we fix *ε* manually to 100 seconds based on the typical time it takes to reset the experimental setup. A prestress was allowed for in the fit to *N* = 2*/*7 control embryos.

## Appendix C Details of the mechanical models

### Overview

In total, we consider four different constitutive models in this article. Written here in one-dimensional form for simplicity, their constitutive equations are (for a stress *σ* and a shear rate 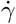):

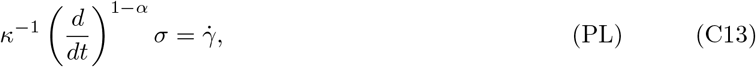

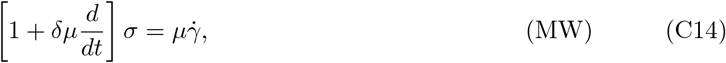

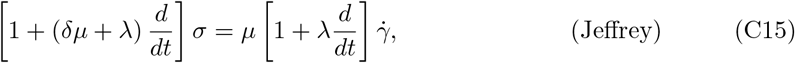

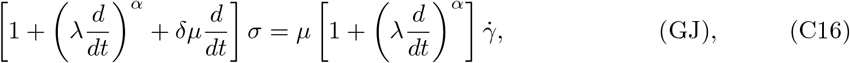

where (*d/dt*)^*α*^ is the fractional derivative operator defined in section D.2. They are the 2-parameter pure power-law (PL) and classical Maxwell (MW) models, the 3-parameter Jeffrey model, and the 4-parameter generalised Jeffrey model (GJ). The GJ, Jeffrey, and Maxwell models form a hierarchy in which each model is obtained as a limiting case of the previous one. Specifically,

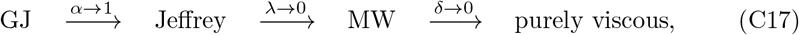

where we also mention purely viscous flow because it was considered in earlier work on this tissue [27]. This hierarchy allows one to think of the models on a scale from more realistic and detailed to more phenomenological and simplified. As discussed in the main text, the additional detail in the more complex models is most prominent on time scales of 𝒪 (100) seconds or less, which is short compared to the time scale of gastrulation which is 𝒪 (10) hours. This explains why the approximation of an embryo as viscous is appropriate in the latter context.

The parameters of these models have the following interpretations:

- *μ*, with units of stress × time, describes a steady-state viscosity
- *δ*, with units of 1*/*stress, is a modulus for the reversible part of the compliance
- *λ* is a time scale for the saturation of this reversible compliance, and
- *α* is a dimensionless exponent of compliance power-law behaviour.

Additionally, the power-law model can be thought of a special case of the GJ model in the triple limit *μ, δ, λ→ ∞* with power-law modulus *κ* = *λ*^*α*^*/δ* constant. It is however otherwise incompatible with the other models, since it does not allow for a steady flow state at long times. The units of *κ* are stress × time^*α*^, which implies that it only has a phenomenological interpretation. Regardless, we can compute *κ* for the other models to compare fits.

Finally, we can define a stress relaxation time Λ = *δμ* + *λ* for the Jeffrey and Maxwell models. For these models, stresses decay exponentially on this time scale. For the GJ and power-law models, stresses decay more slowly as a power-law, in a way that depends also on the history of stress-loading. For this reason, a stress relaxation time scale can not be defined for these models.

The parameters and model relationships are summarised in table C1.

In the following we provide a discussion of each individual model, as well as expressions for its relaxation modulus *G*(*t*) and compliance *J* (*t*) as discussed in section B.

### Power-law model

The power-law model interpolates between elastic (*α* = 0) and viscous (*α* = 1) behaviour. The compliance and relaxation modulus for this model are:

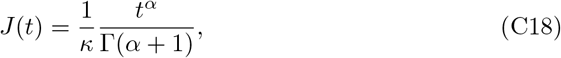

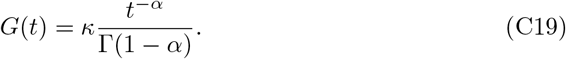

**Table C1.** Parameter relationships between the models. An x indicates an independent parameter. All parameters are strictly non-negative.

| | $\mu$ | $\delta$ | $\lambda$ | $\alpha$ | $\kappa$ | $\Lambda$ |
| --- | --- | --- | --- | --- | --- | --- |
| units | stress ×<br>time | 1/stress | time | – | stress ×<br>time$^\alpha$ | time |
| inter-<br>pretation | steady-state<br>viscosity | compliance<br>modulus | compliance<br>time | power-law<br>exponent | power-law<br>modulus | stress-<br>relaxation<br>time |
| GJ | x | x | x | x | $\lambda^\alpha/\delta$ | undefined |
| Jeffrey | x | x | x | 1 | $\lambda/\delta$ | $\delta\mu + \lambda$ |
| MW | x | x | 0 | – | 0 | $\delta\mu$ |
| PL | ∞ | ∞ | ∞ | x | x | undefined |

The compliance grows as a power-law with exponent *α*. The relaxation modulus decays as a power-law with exponent − *α*. In the elastic limit *α →*0, the compliance is a step of amplitude 1*/κ* and the relaxation modulus is constant (i.e. does not decay). In the viscous limit, compliance is linear while the relaxation approaches a Dirac delta-function. The model is reversible for any value of *α <* 1, in the sense that *γ →*0 if any stresses are released. The time scale on which this happens is determined by the duration for which the stresses were applied. In more detail, we have for *σ* = Θ(*t*) − Θ(*t* − *T*):

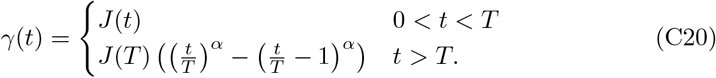

The second parameter *κ* has units of stress × time^*α*^. Since these units are not physical for 0 *< α <* 1, the parameter is phenomenological. It is possible to think of it as a compound parameter that arises from multiplying a viscosity with a time scale to some power, but there is no unique way to define these, or determine them individually from experiments. Equivalently, a finite physical compliance with units of 1*/*stress cannot be expressed purely in terms of the model parameters, and hence depends on the extrinsic time scale on which stresses are applied. The parameter *κ* is therefore best thought of as a scale factor.

### Maxwell model

The Maxwell model is again straightforward to infer from the constitutive equation. Using the definition Λ = *δμ* we have the compliance *J*, and relaxation modulus *G* as

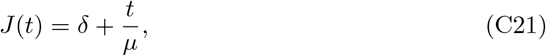

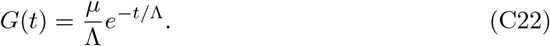

The compliance is a step followed by a linear flow. The viscosity *μ* controls the rate of extension, and *δ* controls the magnitude of reversible elastic compliance in response to stress. The relaxation modulus also decays exponentially on a time scale Λ, which justifies its interpretation as the stress relaxation time scale. In the limits Λ →0 and Λ→ ∞, the model becomes purely viscous (with viscosity *μ*), and purely elastic (with spring constant 1*/δ*) respectively. The model is partially reversible and strains respond instantaneously and discontinuously when stresses are released. For *σ* = Θ(*t*) − Θ(*t − T*):

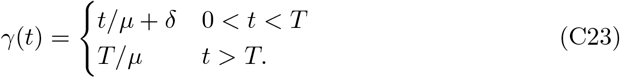

### Jeffrey model

Defining the stress relaxation time Λ = *μδ* + *λ*, we find for this model that

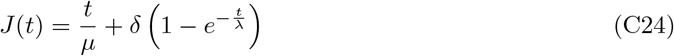

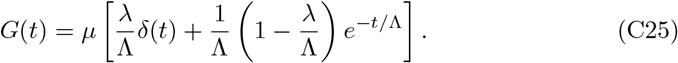

One can think of this model as an extension of the Maxwell model with an additional time scale *λ* that regularises the jump in displacement after a stress is applied. For this reason *λ* called the compliance time. The viscosity *μ* controls the slope of the steady-state flow, and *δ* defines an elastic compliance that is consistent with the other models, and can be thought of as a viscoelastic “buffer” with fill/depletion rate *λ*. The relaxation modulus decays exponentially on a time scale Λ (≠ *λ*), but also contains a Dirac-delta, indicating a viscous behaviour at very short time scales.

Note that for a physically admissible material, with *G*(*t*) decaying monotonically, stress relaxation is always slower than strain retardation Λ ≥ *λ*, and in practice usually Λ ≫ *λ*. The Jeffreys model is partially reversible, with strain decaying exponentially on the same scale *λ* as during loading. We have for *σ* = Θ(*t*) − Θ(*t* − *T*):

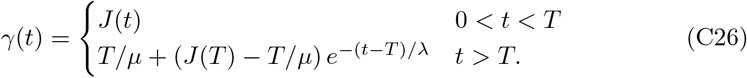

### Generalised Jeffrey model

The Jeffrey model regularises the Maxwell model with a non-zero compliance time scale *λ* on which the flow exponentially approaches a linear creep with viscosity *μ* when under stress. As discussed above, this model predicts equally that the strain relaxes on the same time scale *λ* when the stress is removed. To account for potentially different loading and relaxation time scales, as well as more general than exponentially decaying approaches to the steady state, we generalise the Jeffrey model by promoting the parallel dashpot with viscosity *λ/δ* to a fractional element with modulus *κ* = *λ*^*α*^*/δ*.

This 4-parameter model, which we call the *generalised Jeffrey model (GJ)*, is described by the constitutive equation

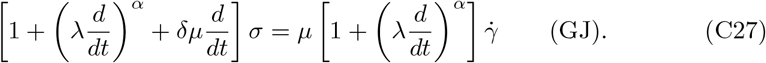

We recover the Jeffrey model is recovered in the limit *α* → 1, and it is natural to think of 1 − *α* as a small parameter that measures the divergence from that model.

This model is very versatile, and can exhibit a very wide range of different behaviours depending on what limits are taken. This results in potentially unrealistic fits that don’t take into account known properties of the epiblast. For this reason, we constrain *μ* ≤ *μ*_max_, enforcing a finite viscosity and the existence of steady-state viscous behaviour, and *δ* ≤ *δ*_max_, ensuring that the reversible part of the compliance is finite.

The Laplace transform of the relaxation modulus for the GJ model is

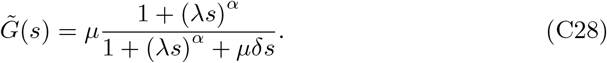

This expression can be written as a series expansion around *s* =∞ and inverted term by term to find the viscoelastic modulus *G*. Unfortunately, the resulting series has very poor numerical convergence for *α* close to 1, which is the limit that we are most interested in.

Instead, it is more convenient to perform a 𝒥-fit of this model. To this end we obtain we the Laplace transform of the 𝒥-operator kernel as

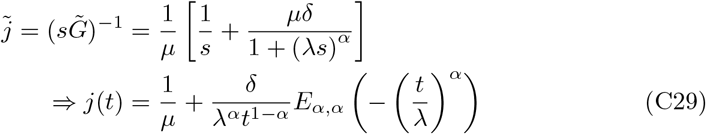

Here the Mittag-Leffler function *E*_*α,β*_(*z*) is defined as

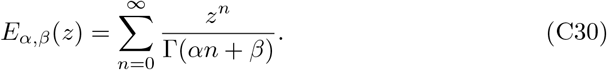

Detailed information on the properties of this function may be found elsewhere [6, 44]. The compliance is then

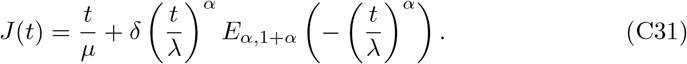

Since *E*_1,2_(*z*) = *z*^−1^ (*e*^*z*^ − 1), we recover the exponential approach to a steady state for the Jeffrey model as *α* → 1. For *α <* 1 we can understand the behaviour of the compliance by considering asymptotic expansions of *E*_*α*,1+*α*_ as *t/λ* →0 or → ∞. This gives

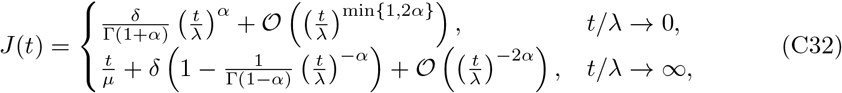

which shows that the reversible compliance initially grows quickly as a *t*^*α*^ power-law and then approaches maximum capacity slowly as a *t*^−*α*^ power-law, with *λ* indicating the cross-over time scale between the two behaviours. Note that when *α* significantly differs from 1, *λ* can grow very large without a significant increase of the modulus *κ* = *λ*^*α*^*/δ*. In practice this means that *κ* is a more suited as a fitting parameter despite lacking an immediate physical interpretation.

## Appendix D Mathematical definitions

### D.1 Heaviside steps and Dirac deltas

We refer to the Heaviside step function as

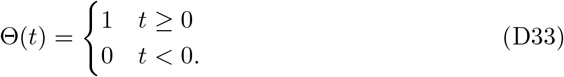

We also employ the Dirac delta distribution *δ*(*t*) that is defined by the sampling property,

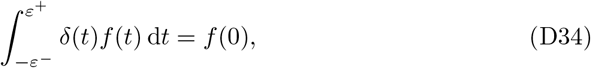

for an arbitrary function *f* and *ε*^+,−^ *>* 0. The integral is in general not well-defined if either of the integration limits is exactly 0. In this article however we tacitly consider Eq. (D34) to hold also when either of *ε*^+,−^ = 0. Physically, our integration variable represents time, and with this convention the causal effect of the Dirac delta takes place fully within the considered time interval.

### D.2 Fractional integrals and derivatives

In this paper we use the Caputo definition of the fractional integral and derivative as follows [44, 45]:

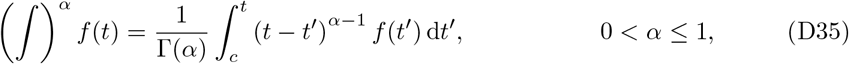

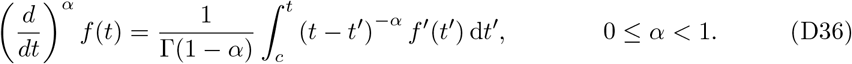

Here we note that (*d/dt*)^*α*^ ≡ (∫)^1−*α*^(*d/dt*). *A priori* these definitions are valid only for *t* ≥ *c*, where *c* is an undetermined constant. To see its effect, we can substitute *α* = 0 in Eq. (D36) to find

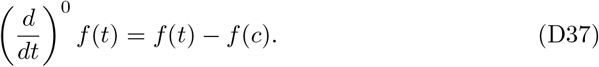

Thus, in order for the definition to make sense we require *f* (*c*) = 0. If we restrict ourselves to functions *f* with *f* (*t*) = 0 for *t* ≤ *c*, then the definitions become universally valid for any value of *t*. Without loss of generality then we can choose *c* = 0^−^ for an experiment started from an unperturbed state at *t* = 0^−^. With this convention we have the Laplace transforms

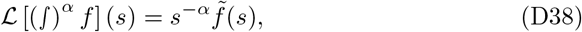

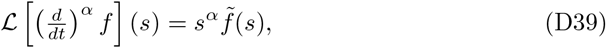

where we use the notation 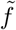 to denote the Laplace transform of *f*.

### D.3 Integral operators and kernels

A general linear viscoelastic model can be defined by its relaxation modulus *G*(*t*), which characterises the stress response to a step strain. Mathematically, this gives rise to an integral operator 𝒢 that acts on the strain rate 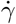 :

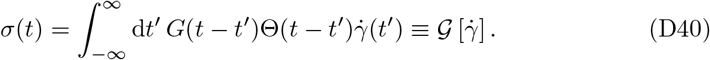

With the conventions noted in section D.2, we then have *G*(*t*) = 𝒢 [*δ*(*t*)].

For physically realisable models, this operator is invertible. The inverse, which describes the kinematic response to an imposed stress, can be more intuitive to think about, and is defined as follows:

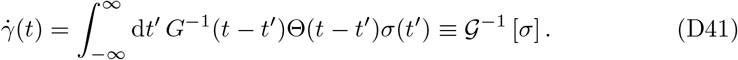

We can integrate once more to obtain the strain, and define the compliance operator 𝒥:

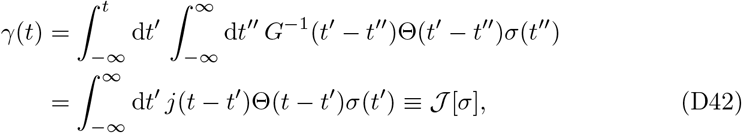

where 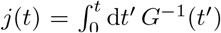. The compliance *J* (*t*) is defined as the strain response to a unit step stress, *J* (*t*) = 𝒥[Θ(*t*)].

The operators 𝒢, 𝒢^−1^ and 𝒥all describe convolutions of stress and strain with a kernel. Their relationships are therefore expressed conveniently in the Laplace domain:

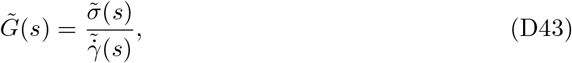

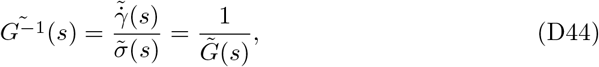

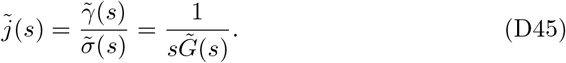

